# Characterization of an enhancer element that regulates *FOXP2* expression in human cerebellum

**DOI:** 10.1101/641597

**Authors:** Julen Madurga, Francisco J. Novo

**Affiliations:** Department of Biochemistry and Genetics. School of Science. University of Navarra, 31008 Pamplona (Spain)

## Abstract

Human cerebellum is involved not only in motor control but also in several cognitive skills. In this work, our goal was to identify genomic enhancers responsible for the regulation of genes implicated in the development of this brain region. From an initial collection of Vista elements showing specific enhancer activity in mouse hindbrain, we found an enhancer element located within the final intron of *FOXP2*, a gene expressed in cerebellum and implicated in vocal communication and speech and language disorders in humans. Analysis of this enhancer using various computational resources suggests that it is a strong candidate to account for *FOXP2* expression during cerebellar development. Two blocks within this region are deeply conserved in vertebrates; they are separated by a microsatellite sequence present only in eutherian mammals, and expanded in human genomes.

## Introduction

The cerebellum is a good model for the study of neurogenesis, neurodevelopment and brain evolution (1). It contains about 80% of the neurons of the CNS, even though it only represents 10% of its total mass (2). As well as its involvement in motor function, the cerebellum also participates in cognition, and shows a high degree of connectivity with the neocortex (3). In keeping with its involvement in cognitive functions, the cerebellum has expanded in hominid evolution, even when comparing archaic and modern humans (3). Likewise, cerebellar syndromes in humans frequently display some type of cognitive impairment.

As reviewed by *Carroll* (4), the suggestion by King and Wilson that phenotypic differences between human and chimpanzee would mostly result from regulatory changes has been subsequently confirmed; numerous findings have shown changes in cis-regulatory elements during human evolution, of which those affecting brain development are particularly relevant. *FOXP2* is a good example of how changes in regulatory elements could have altered gene expression patterns during brain development and thus drive the evolution of cognitive abilities such as vocal learning. Recent research has shown the importance of transcriptomic changes during brain development in primate evolution (5,6), and the role of enhancers in orchestrating such changes: many enhancers regulating genes involved in brain development harbour human-specific variants, which are frequently associated with neurodevelopmental problems (7).

As the role of enhancers in cerebellum evolution has not been extensively studied, in this work we have used several computational tools with a view to identify enhancers that are active in this brain region and regulate genes expressed during development. We have found a previously described enhancer element for which chromatin marks, gene expression profiles and chromatin contact data collectively suggest a role in *FOXP2* regulation during cerebellar development.

## Methods

We used several datasets available as tracks or track hubs at the UCSC Genome Browser. In order to identify and analyse enhancer regions, we took annotations from *Vista enhancers* (8) and *GeneHancer* (9). Enhancer activity in neural tissues was predicted using annotations from *IDEAS* (10,11) to visualize chromatin states (derived from chromatin marks) in neural tissues. Transcription factor binding sites (TFBS) from ChIP-seq data were obtained from *Remap* (12), which includes 31 million ChIP-seq peaks from ENCODE and 49 million peaks from publicly available data.

*Gene expression-GTEx RNA seq* (13) was used to analyse gene expression in adult tissues. *BrainSpan* data (14) for specific genes were viewed in *GenTree* (15); the *Human Developmental Biology Resource* (16) was accessed via *Expression Atlas* (17). Both resources were used to analyse gene expression in neural tissues during various developmental stages.

*GTEx Tissue eQTL* (18), *ClinVar Short Variants* (19) and *GWAS Catalog* (20) were inspected to search for association between gene expression or neuro-developmental phenotypes with specific variants located inside enhancers.

*Enrichr* (21,22) was used for enrichment analysis of biological categories (gene ontologies, biological pathways and expression levels in different tissues) in gene lists.

Physical interactions between candidate regulatory elements and surrounding gene promoters were assessed as virtual 4C plots either in the *3D Genome Browser* (23) or in *3DIV* (24), comparing virtual 4C plots in several neural and non-neural tissues.

The conservation track in the UCSC Genome Browser showed Multiz alignments in selected vertebrate genomes. Further alignments of candidate regulatory regions with genomic sequences were performed with *CoGe Blast* (25). Each candidate regulatory element was inspected in detail, selecting the core region more likely responsible for enhancer activity (PhastCons score, epigenetic marks and presence of TFBS) and used this as query in blastn searches against the genomes of species representing a range of vertebrate taxons. Table 1 lists the genome assemblies that were used in these searches.

**Table 1.** Genome assemblies used in *COGE* blastn searches.

| Species | Genome assembly |
| --- | --- |
| *Homo sapiens* | vGRCh37 (id 29349) |
| *Pan troglodytes* | v2.1.4 (id28817) |
| *Otolemur garnettii* | v3 (id 16664) |
| *Mus musculus* | vGRCm38.p5 (id 34107) |
| *Bos taurus* | v3.1 (id 34644) |
| *Loxodonta africana* | v3 (id 16685) |
| *Monodelphis domestica* | v5 (id 35487) |
| *Ornithorhynchus anatinus* | v5.0.1 (id 28045) |
| *Gallus gallus* | v4 (id 28417) |
| *Xenopus tropicalis* | v9.1 (id 52485) |
| *Latimeria chalumnae* | v1.0 (id25005) |
| *Danio rerio* | v10 (id 25001)0 |
| *Gasterosteus aculeatus* | v2 (id 52353) |
| *Takifugu rubripes* | v5 (id 25444) |
| *Tetraodon nigroviridis* | vUCSC (id23254) |
| *Lepisosteus oculatus* | v1 (id 52331) |
| *Callorhynchus milii* | v6.1.3 (id 25133) |
| *Eptatretus burgeri* | v3.2 (id 52454) |
| *Petromyzon marinus* | v7.0.64 (id 12390) |

## Results

We searched for all VISTA elements that show activity in mouse hindbrain, either exclusively or in hindbrain and neural tube. This resulted in a set of 58 enhancers (38 active exclusively in hindbrain and 20 active in hindbrain and neural tube). In order to confirm the reliability of this dataset, we made a list with all genes flanking these enhancers up to a distance of 1 Mb in each direction (93 genes) and analysed the enrichment of several biological categories using Enrichr. As shown in Table 2, this gene list is significantly over-represented (p_adjusted_<0.05) in genes upregulated in several brain regions according to *ARCHS*^*4*^ *Tissues* database (26), cerebellum showing the highest absolute Z-score value (1.69). A similar result was obtained in the *Cancer cell line Encyclopedia* database, with a significant enrichment in genes expressed in neural cell lines (p_adjusted_=2.1×10^-4^); in the *Tissue Protein Expression from Human Proteome Map* database, significant enrichment was found exclusively in fetal brain, (p_adjusted_=0.014), and in *Jensen Tissues* database (27) the highest absolute Z-score (4.12) corresponded to *ectoderm* (p_adjusted_=5.3×10^-4^).

**Table 2.** Enrichment analysis of 93 genes located near Vista elements active in hindrain. Genes in this list are enriched in genes upregulated in brain regions, according to *ARCHS*^*4*^ *Tissues* database. Only significant values (adjusted p-value) are shown. Z-scores indicate, for every category, the variation in rank (standard deviations) in the gene list compared to the background list.

| Name | P-value | Adjusted p-value | Z-score | Combined score |
| --- | --- | --- | --- | --- |
| NEURONAL EPITHELIUM | 3.508e-7 | 0.00003789 | -1.58 | 23.52 |
| PREFRONTAL CORTEX | 0.00001297 | 0.0007006 | -1.54 | 17.36 |
| CEREBELLUM | 0.00003909 | 0.001055 | -1.69 | 17.13 |
| MOTOR NEURON | 0.00003909 | 0.001055 | -1.64 | 16.66 |
| SPINAL CORD | 0.0001118 | 0.002013 | -1.62 | 14.75 |
| SPINAL CORD (BULK) | 0.0001118 | 0.002013 | -1.59 | 14.46 |
| MIDBRAIN | 0.0007797 | 0.009357 | -1.46 | 10.45 |
| RENAL CORTEX | 0.0007797 | 0.009357 | -1.43 | 10.23 |
| FETAL BRAIN CORTEX | 0.0007797 | 0.009357 | -1.39 | 9.97 |
| FETAL BRAIN | 0.001897 | 0.01707 | -1.47 | 9.23 |

These results support the assumption that these Vista enhancers collectively tend to regulate genes involved in neurodevelopmental processes. In order to identify which of these enhancers might be associated with cerebellum, we used a combination of approaches: (i) presence of eQTLs or variants associated with a neurodevelopmental phenotype (by GWAS or in ClinVar); (ii) expression pattern of the genes surrounding each Vista element should be consistent with its involvement in cerebellar function, and (iii) presence of epigenetic marks of active enhancer in neural tissues. This led to a short list of three Vista enhancers very likely to be involved in cerebellar function and/or development, shown in Table 3.

**Table 3.** VISTA enhancers selected after initial analysis. “r” indicates that the enhancer is active in rhombencephalon, “nt” in neural tube. Genome coordinates in GRCh37 assembly.

| Enhancer | GRCh37 coordinates | Active | Epigenetic marks of active enhancer in: | Overlaps with: |
| --- | --- | --- | --- | --- |
| hs966 | chr7:114326912-114329772 | r | Neural (embryonic and adult) | FOXP2 |
| hs1082 | chr11:31816452-31818421 | r + nt | Neural (embryonic) | PAX6 |
| hs2094 | chr1:10795106-10799241 | r | Neural and non-neural | CASZ1 |

The hs966 Vista element is located in the final intron of *FOXP2*, shows high and specific activity in mouse hindbrain according to Vista and bears marks of active enhancer in several brain samples and progenitor cell lines from the *Roadmap Epigenetics Project* (Figure 1): germinal matrix (E070), male fetal brain (E081), female fetal brain (E082), H1 ESCs (E003) and H9 derived neuronal progenitor cells (E009).

**Figure 1.**
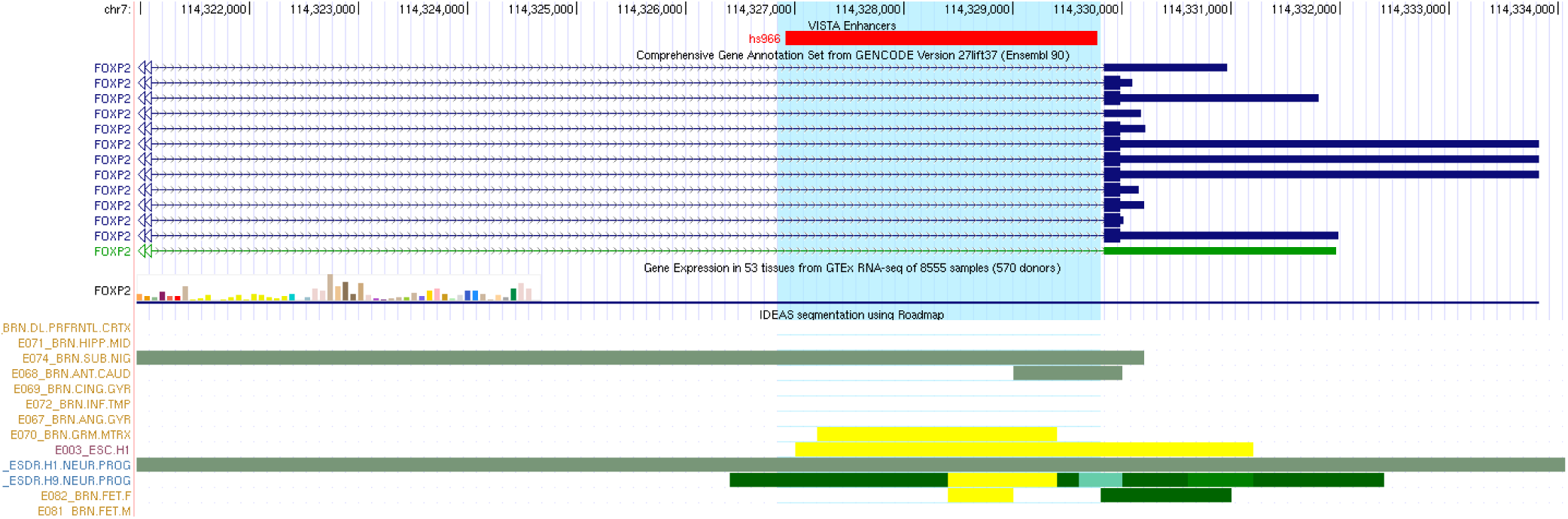
Location and chromatin marks of Vista element hs966 (in red at the top). Rectangles in the bottom tracks indicate active enhancer status (yellow) or transcription (green) according to the IDEAS Roadmap chromatin segmentation method.

As all Vista elements, hs966 is included in *GeneHancer* database (enhancer GH07J114686), which predicts its association with *FOXP2* only on the grounds of its distance to the promoter of this gene (no evidence based on eQTLs, capture HiC, eRNA co-expression or transcription factor co-expression). Given the prior association of *FOXP2* with vocal communication through Purkinje cell development and with speech and language disorders (28-30) we decided to search for further evidence that the hs966 element constitutes a good candidate to regulate *FOXP2* expression in cerebellum during development.

First, we searched for data on *FOXP2* expression throughout human development. Although expression in adult brain tissues is not particularly high according to GTEx, data from *Brainspan* show a clear peak of *FOXP2* expression in cerebellum, and to a lesser extent in midbrain, around post-conception week (pcw) 12 of human embryonic development (Figure 2). Expression goes down quickly and is then maintained at lower levels until 32 pcw when it decreases to postnatal levels. A lower expression burst is also evident in embryonic striatum, again disappearing in late-prenatal and postnatal samples.

**Figure 2.**
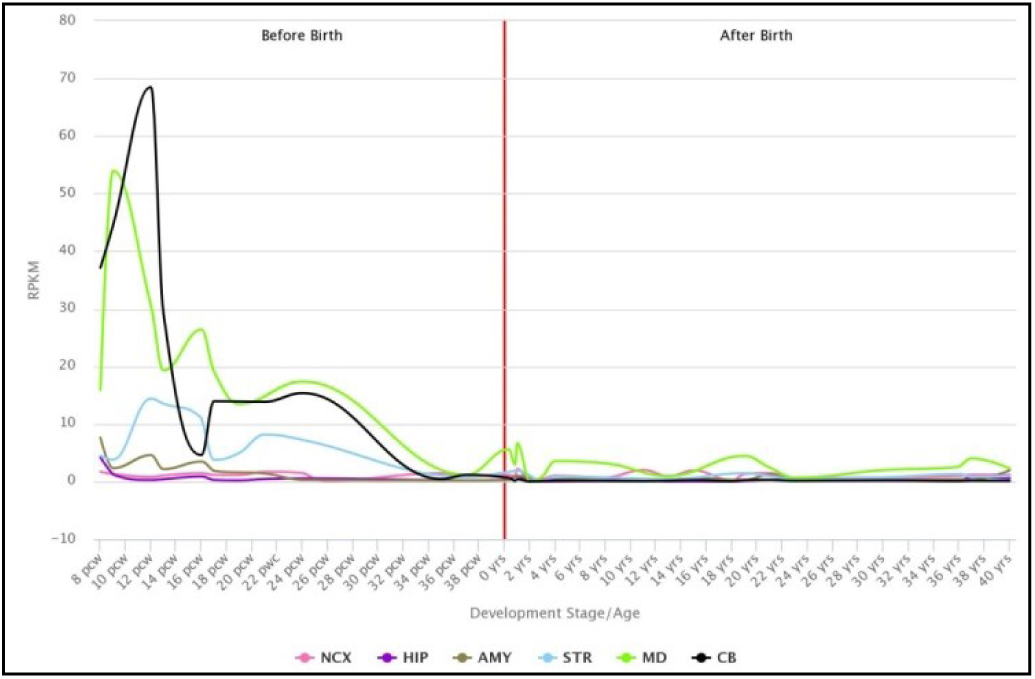
*FOXP2* expression during development and postnatal life in six different brain regions according to data from *Brainspan*. The black line shows the expression profile in cerebellum, which peaks at 12 pcw (post conception week).

Next, we searched for HiC datasets from brain samples and compared them with non-neural tissues, in an effort to find further experimental evidence for the contact between this enhancer and the promoter of *FOXP2*. Although, to the best of our knowledge, HiC experiments are not available in human developing cerebellum or Purkinje cells, we explored contacts in datasets from H1 neuronal progenitor cells (GSM1267202; GSM1267203), cerebellar astrocytes (ENCLB672PAB; ENCLB174TEA), hippocampus (GSM2322543) and H1 embryonic stem cells (GSM1267196; GSM1267197), as well as aorta (GSM1419084) and lung (GSM2322544; GSM2322545). Virtual 4C plots revealed a clear interaction between the hs966 element and the major promoter of *FOXP2* in aorta, neural progenitors (shown in Figure 3) and hippocampus, but not in stem cells or astrocytes from cerebellum.

**Figure 3.**
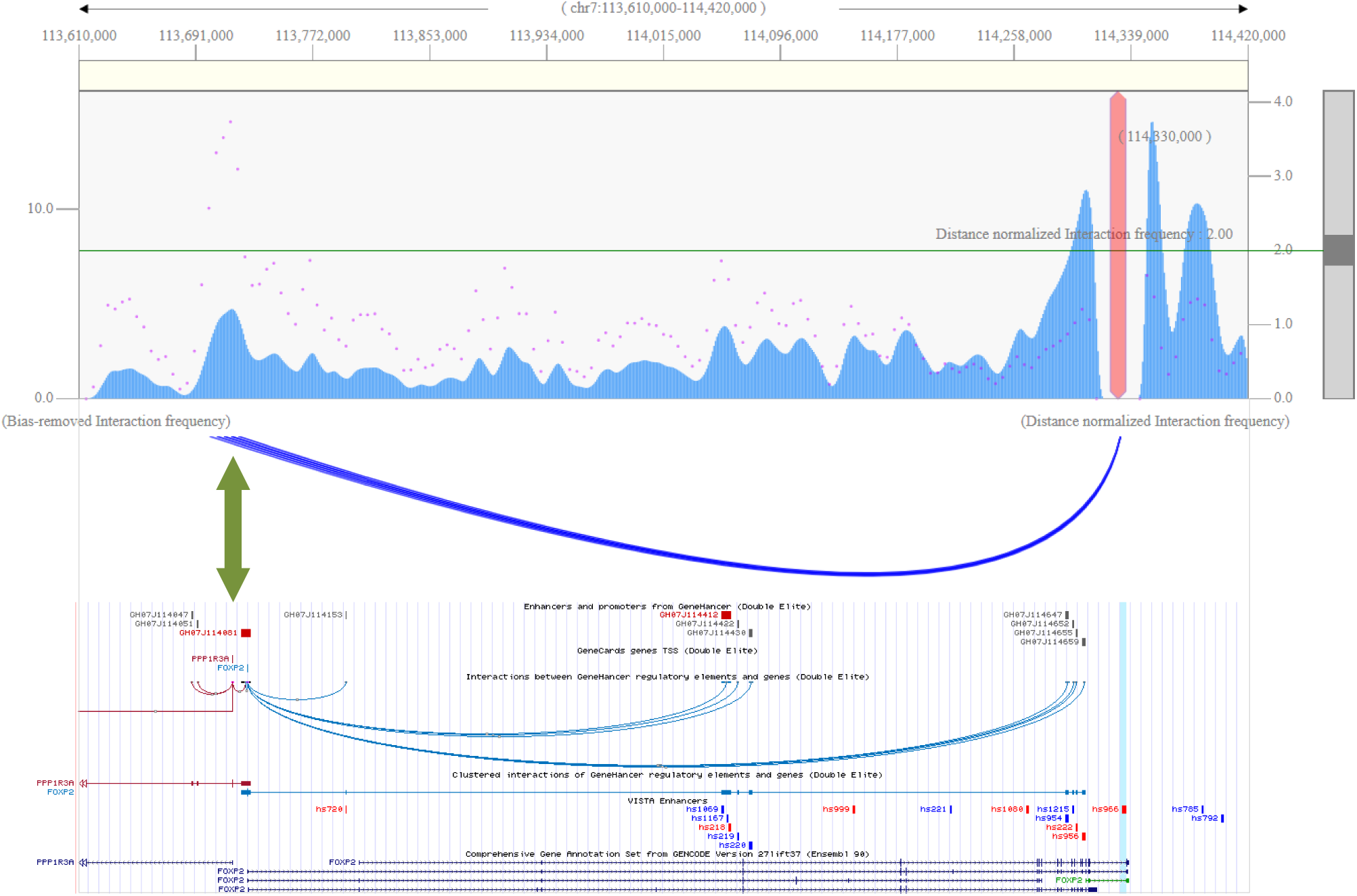
Top: virtual 4C plot showing number of contacts (pink dots) and contacts with distance-normalized interaction frequency>2.0 (blue arches) between Vista enhancer hs966 (red vertical bar) and the major promoter of *FOXP2* (thick green arrow) in neuronal progenitors using the 3DIV tool. The same region is shown in the UCSC Genome Browser at the bottom, including Gencode annotations, GeneHancer elements and Vista enhancers (hs966 highlighted in blue). Coordinates correspond to the GRCh37 assembly.

Taken together, both the expression profile of *FOXP2* during brain development and chromatin interaction data between its promoter and the hs966 Vista element support a role for this enhancer in regulating the expression of this gene in embryonic brain.

Conservation of the hs966 element also supports its functional relevance. As depicted in Figure 4, Multiz alignments in 100 vertebrate genomes reveal very deep conservation. Within this 2.8 kb element, two smaller blocks show greater constraint with high PhastCons scores. These two regions contain binding sites for several transcription factors in ChIP-seq experiments from neural and non-neural samples (Figure 4, top), as well as a peak of p300 binding in neural cells originated from H1-hESC (ENCSR843ZUP). Both are well conserved in sarcopterygians and actinopyerygians, but not in lamprey.

**Figure 4.**
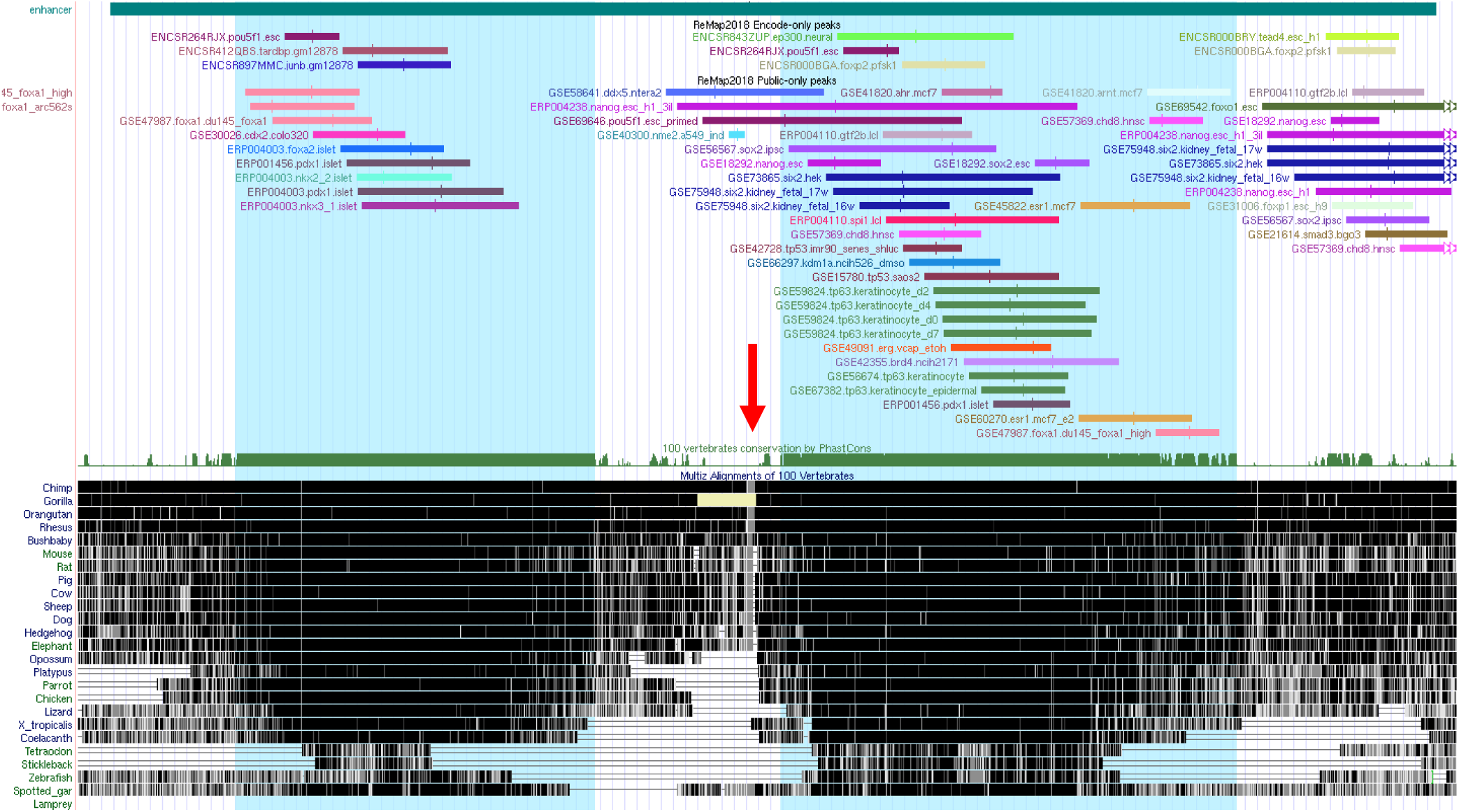
Evolutionary analysis of hs966 Vista element. Inside this 2.8 kb element (green rectangle at the top) two smaller regions of great conservation are evident (highlighted in blue), with high PhastCons scores. Conservation in representative vertebrate taxons is shown at the bottom. Transcription factor binding sites from *Remap* (ENCODE and other publicly available ChIP-seq experiments) are also shown. Both highlighted regions show several TFBS peaks in various cell lines. A region of low conservation (red arrow) includes a microsatellite which is polymorphic in humans but is absent in most vertebrate genomes (see text).

We then used the sequence of these blocks as query in blastn searches using COGE blast. We found significant hits in the genomes of actinopterygians and in the elephant shark, but not in lamprey or hagfish. Interestingly, alignments were much more significant in spotted gar and elephant shark than in teleostei genomes (which included *Danio rerio, Gasterosteus aculeatus, Tetraodon nigroviridis* and *Takifugu rubripes*). For instance, blastn hits for the left block had 73% coverage of the query sequence and E-value=0.0 in the spotted gar genome, whereas coverage of the query varied between 7.2% (E-value=6e-04) to 21.2% (E-value=2e-21) for *Tetraodon* and *Danio*, respectively, and was completely absent in *Takifugu*. A similar pattern was found for the right block, with very significant hits in *Callorhinchus* and *Lepisosteus* (85% coverage of the query sequence and E-value=0.0 in both cases) compared to teleostei (from 23.1% coverage and E-value=8e-25 in *Danio* to 22.7% and E-value=2e-33 in T*akifugu*). In fact, all hits in teleostei genomes were fragmented in two or three blocks.

Of note, the region between these two highly-conserved blocks shows little conservation, particularly a 90 bp segment which seems to be absent even in great ape genomes. Closer inspection of this segment revealed the presence of an (AC)_19_ tandem repeat, flanked by two T-rich low complexity tracts. This microsatellite is very polymorphic in humans (it spans several SNVs from dbSNP) and is also present in the high-resolution denisovan genome. Primates have a shorter version of 8 repeats, also followed by a poly(T) stretch; however, euarchontoglires, laurasiatheria, and afrotheria have the short version of the dinucleotide repeat without the poly(T) tract, whereas the whole segment is absent in non-eutherian mammals and more basal vertebrates.

## Discussion

*FOXP2* is expressed in mouse and human developing cerebellum and it has been previously implicated in speech and language disorders (31,32). It might be crucial for Purkinje cell development, as it is specifically expressed in those cells, and thus play a relevant role in vocal communication (28-30). Our searches in recent neurodevelopmental expression databases confirmed that *FOXP2* expression in developing human brain is highest in cerebellum, around 12 pcw. However, there is so far no evidence as to the identity of the cis-regulatory elements responsible for cerebellum-specific expression of this gene.

Our search for Vista enhancers with activity in hindbrain led us to hs966, an enhancer that shows strong and specific enhancer activity in mouse rhombencephalon in embryonic day e11.5, in 6 out of 11 embryos analysed by the Vista enhancer project. This 2.8 kb element is located in the final intron of *FOXP2*, suggesting that it might regulate this gene, but experimental evidence for this association was lacking. In our work we have found that Vista element hs966 displays chromatin marks of active enhancer in human fetal brain samples and in neuronal progenitor cells and that it contacts the major *FOXP2* promoter in H1-derived neuronal progenitor cells. The presence of ChIP-seq peaks for several transcription factors in neural and non-neural samples reinforces the role of hs966 as an enhancer. Therefore, we propose that this element is a strong candidate to account for cerebellum-specific expression of *FOXP2* during development.

As most Vista elements and developmental enhancers, hs966 is deeply conserved in vertebrate genomes. Two blocks, in particular, are very well conserved in all gnathostome genomes analysed but not in agnatha (lamprey and hagfish). However, conservation of these blocks is weaker in the genomes of teleostei. This suggests that the core elements of the hs966 enhancer were present in the gnathostome ancestor and were conserved in sarcopterygians and non-teleostean actinopterygiians.

Interestingly, these two blocks are separated by a short segment formed by a microsatellite, which is present only in eutherian mammals. Whereas primates and mammals have shorter versions, the microsatellite is expanded in the genomes of anatomically modern humans and denisovans, showing a high degree of polymorphism in repeat number in modern humans. Differences in the size of cerebellum between modern humans and neandertals have been suggested to underlie the cognitive differences that allowed the development of more sophisticated tool usage, language and social skills in modern *Homo sapiens* (34). It is tempting to speculate that the expansion of this microsatellite in the middle of hs966 might have optimized the function of this enhancer in humans, and thus increased its regulatory activity on the promoter of *FOXP2* during development of the cerebellum.

Several lines of research should be undertaken to confirm our prediction. Deletion of this Vista element in model vertebrates will show whether it plays a significant role on *FOXP2* expression in cerebellum at different stages of embryonic development, and any potential structural of functional abnormality derived from its absence. This should be complemented with single-cell transcriptomic analysis of Purkinje and progenitor cells in model vertebrates, as well as chromatin conformation studies aimed at confirming the contact between this enhancer and the promoter of *FOXP2* in the same samples.

## Acknowledgments

We would like to thank Dr. José L. Vizmanos for helpful discussions.

